# Acyl-ghrelin attenuates neurochemical and motor deficits in the 6-OHDA model of Parkinson’s disease

**DOI:** 10.1101/2022.01.31.478447

**Authors:** Daniel Rees, Amy L. Johnson, Mariah Lelos, Gaynor Smith, Luke D. Roberts, Lindsay Phelps, Stephen B. Dunnett, Alwena H Morgan, Rowan M Brown, Timothy Wells, Jeffrey S. Davies

**Author notes:** Corresponding author, Address for editorial correspondence: Dr Jeffrey S Davies, Molecular Neurobiology, Institute of Life Science, School of Medicine, Swansea University, UK. SA28PP.

## Abstract

The feeding-related hormone, acyl-ghrelin, protects dopamine neurons in murine 1-methyl-4-phenyl-1,2,3,6-tetrahydropyridine (MPTP)-based models of experimental Parkinson’s disease (PD). However, the potential protective effect of acyl-ghrelin on substantia nigra pars compacta (SNpc) dopaminergic neurones and consequent behavioural correlates in the more widely used 6-hydroxydopamine (6-OHDA) rat medial forebrain bundle (MFB) lesion model of PD are unknown. To address this question, acyl-ghrelin levels were raised directly by mini-pump infusion for 7-days prior to unilateral injection of 6-OHDA into the MFB with assessment of amphetamine–induced rotations on days 27 and 35, and immunohistochemical analysis of dopaminergic neurone survival. While acyl-ghrelin treatment was insufficient to elevate food intake or body weight, it attenuated amphetamine-induced circling behaviour and SNpc dopamine neurone loss induced by 6-OHDA. These data support the notion that elevating circulating acyl-ghrelin may be a valuable approach to slow or impair progression of neurone loss in PD.

**Highlights:**

- Acyl-ghrelin attenuates SNpc dopamine cell loss in rat 6-OHDA-lesion model of PD
- Acyl-ghrelin attenuates motor deficits in rat 6-OHDA-lesion model of PD

## Introduction

Parkinson’s disease (PD) is the second most common neurodegenerative disorder affecting an estimated 6.1 million people worldwide (GBD 2016). The disease is characterised by age-related loss of dopamine-containing neurones in the substantia nigra *pars compacta* (SNpc) region of the adult brain. Alpha-synuclein immunopositive aggregates, termed Lewy bodies, are a neuropathological hallmark of PD that form in neurones (1). This neuroanatomical pathology results in the loss of motor (bradykinesia, resting tremor) and non-motor (cognitive decline, sleep disturbance, constipation) function. Current therapies are available to control symptoms of disease, however, a large proportion of patients do not respond to treatment. In addition, the prolonged use of the dopamine replacement therapy, L-DOPA, often results in debilitating dyskinesias (2). Therefore, there is a major un-met clinical need for new protective and restorative treatments for PD.

Recently identified mutations in familial PD (LRRK2, GLB) and neurotoxins that induce Parkinsonian characteristics in animals converge on the cells’ energy producing organelles, the mitochondria. Parallel studies have identified a clear association between dopamine neurone dysfunction within the SNpc of PD brain and metabolic dysfunction (3). This knowledge has led to the search for molecules that improve mitochondrial function and biogenesis with the aim of enhancing cellular resilience to periods of bioenergetics stress. One such environmental factor that enhances both mitochondrial function and resilience against neurodegeneration is calorie restriction (4), but until recently, the underlying mechanism was not well understood. The stomach hormone, acyl-ghrelin, which is elevated by CR (5) and increases mitochondrial activity in neurones (6), is essential for mediating the beneficial effects of CR in the MPTP-model of PD (7). Acyl-ghrelin signalling induces phosphorylation of the cellular energy sensor, adenosine monophosphate-activated protein kinase (AMPK), to promote cell survival. The selective ablation of AMPK from dopaminergic neurones prevented acyl-ghrelin from protecting against MPTP-induced neurotoxicity. These findings are consistent with reports that treatment with exogenous acyl-ghrelin or synthetic ligands for the ghrelin receptor (GHS-R) protect against MPTP-induced SNpc lesions in mice (8–10).

These effects are thought to be mediated by a direct action of ghrelin on the dopamine containing neurones within the SNpc that express GHS-R (9). In support of this concept, pretreatment with ghrelin attenuated cell loss in MPTP-treated mesencephalic derived cultures (MES2.5 cells) (11). Despite these findings, the potential protective influence of ghrelin in the more widely studied unilateral medial forebrain bundle (MFB) 6-OHDA rat model of PD remain unknown. In this study, we report the efficacy of exogenous acyl-ghrelin treatment in protecting against the neurochemical and behavioural effects of 6-OHDA-induced dopaminergic cell loss in the SNpc.

## Materials and Methods

All animal experiments were conducted under the authority of the Animals (Scientific Procedures) Act, 1986 (UK) and in accordance with the ARRIVE guidelines and were specifically approved by the Cardiff University Animal Welfare Ethical Review Body (AWERB). Male SD rats (weighing 194-241g) were purchased from Charles River (Margate, UK) and housed under conditions of 12h light/12h dark (lights on at 06.00h) with water available *ad libitum.* Rats were fed crushed diet (SDS 801066 RM3(E); 3.64kcal/g) available *ad libitum.* Food intake and body weight were monitored daily. On day 0 rats were prepared with sub-cutaneous osmotic minipumps (Alzet model 2001) primed to deliver acyl-ghrelin (Phoenix Pharmaceucitals; 80μg/24μl/day) for 7 days.

After 1 week on these treatment regimes, rats received an injection of desipramine/pargyline (25mg/kg; 50mg/kg; i.p), were anaesthetised with isolfluorane and mounted in a Kopf stereotaxic frame. The skull was exposed and a burr hole was drilled to introduce a syringe for a single unilateral MFB injection of either vehicle (0.2mg/ml ascorbic acid in 0.9% sterile saline; pH5.0) or 6-OHDA (hydrobromide salt (Sigma Chemicals, UK); 3μg/μl in vehicle). To minimize variability due to degradation of the toxin, the 6-OHDA solutions were freshly made, kept on ice and protected from exposure to light. The solution was injected into the right MFB using the following co-ordinates (relative to bregma) Lateral −1.3; Anterior −4.0 and Ventral 7.0. A total volume of 3μl 6-OHDA was injected at a flow rate of 1μl/min, the syringe was left in place for a further 3 min and then slowly removed. The Sham group underwent the identical surgical procedure except for receiving saline rather than 6-OHDA toxin. The rats received an injection of analgesic (Metacam, 150μg/s.c) after the skin was sutured and they were allowed to recover before being returned to their home caging.

Rats were subjected to rotational testing on days 27 and 35. On the day of testing rats received an injection of methamphetamine (2.5mg/ml/kg/i.p in sterile saline) and immediately placed into individual rotometer bowls. Contralateral and ipsilateral rotations were recorded using Rotorat software for 90 mins.

After rotational testing on day 35 rats were terminally anaesthetised with sodium pentobarbital (200mg/ml) (Euthetal, Merial) until total loss of reflex, nose-anus length measured, and subjected to thoracotomy. Rats were killed by transcardial perfusion of 1.5% paraformaldehyde (PFA). After perfusion, brains were excised and post-fixed by immersion in 1.5% PFA in 0.1M PBS overnight at 4°C. Brains were subsequently cryo-protected by sinking in 30% sucrose at 4°C until further processing. In addition, pituitaries were dissected and weighed, and right femori and tibiae dissected for quantification of bone length using a handheld micrometer.

### Immunohistochemistry

Coronal sections (30μm) were cut along the entire rostro-caudal extent of the mid-brain using a freezing-stage microtome (MicroM, Thermo) and collected (1:12) for free-floating immunohistochemistry. For DAB-immunohistochemical analysis of TH labelling, sections were washed in 0.1M PBS (2×10mins) and 0.1M PBS-T (1×10mins). Subsequently, endogenous peroxidases were quenched by washing in a PBS plus 1.5% H_2_O_2_ solution for 20 minutes. Sections were washed again (as above) and incubated in 5% NDS in PBS-T for 1h. Sections were incubated overnight at 4°C with rabbit anti-tyrosine hydroxylase (1:1000 Millipore, USA) in PBS-T and 2% NDS solution. Another wash step followed prior to incubation with biotinylated donkey anti-rabbit (1:400 Vectorlabs, USA) in PBS-T for 70 minutes. The sections were washed and incubated in ABC (Vectorlabs, USA) solution for 90 minutes in the dark prior to another two washes in PBS, and incubation with 0.1M sodium acetate pH6 for 10 minutes. Immunoreactivity was developed in nickel enhanced DAB solution followed by two washes in PBS. Sections were mounted onto superfrost+ slides (VWR, France) and allowed to dry overnight before being de-hydrated and de-lipified in increasing concentrations of ethanol. Finally, sections were incubated in histoclear (2 x 3 minutes; National Diagnostics, USA) and coverslipped using Entellan mounting medium (Merck, USA). Slides were allowed to dry overnight prior to imaging.

### Imaging and quantification

A one-in-twelve series of 30μm sections (360μm apart) from each animal was immunohistologically stained (see above) and imaged using a light microscope (Nikon Eclipse 50i; 10× objective) and analysed using Image J software. TH^+^ immunoreactive cells were counted bilaterally throughout the entire rostro-caudal extent of the SNpc. Resulting numbers were divided by the number of coronal sections analyzed and multiplied by the distance between each section to obtain an estimate of the number of cells per SNpc. TH ^+^ immunoreactive cells from the lesioned SNpc were expressed as a percentage of the TH^+^ immunoreactive cells present in the un-lesioned SNpc. All experiments were performed in a blinded manner.

For the histological analyses, statistical analyses were carried out using GraphPad Prism 8.0. Statistical significance was assessed by unpaired two-tailed Student’s t-test or one-way ANOVA with Tukey’s post-hoc test for data with normal distribution. Kruskal-Wallis testing with Dunn’s post-hoc analysis was used for data that was not distributed normally: *, *P*<0.05; **, *P*<0.01; ***, *P*<0.001.

## Results

### Acyl-ghrelin treatment did not increase body weight or food intake

Apart from the transient reduction in food intake seen in all groups on the day of lesion surgery, daily food intake (Fig 1A) was not further affected by either the 6-OHDA lesion, nor the 7-day infusion of acyl-ghrelin. However, cumulative food intake was significantly reduced by 6-OHDA lesion (*P*<0.01), an effect that was partially alleviated by pre-treatment with acyl-ghrelin (Fig 1B). Similarly, overall body weight gain was reduced by 20% in rats receiving the 6-OHDA lesion alone (*P*<0.01; Fig 1D), which was also partially alleviated by pre-treatment with acyl-ghrelin. In addition, neither the 6-OHDA lesion, nor acyl-ghrelin pre-treatment had any significant effect on either of the markers of skeletal growth; nose-anus length (Fig 1E), femoral length (Fig 1F) and tibial length (Fig1G). Pituitary weight was not affected by either treatment (Fig1H). Thus, the dose of acyl-ghrelin given was insufficient to induce orexigenesis or amplify the activity of the hypothalamo-pituitary-growth axis.

**Figure 1.**
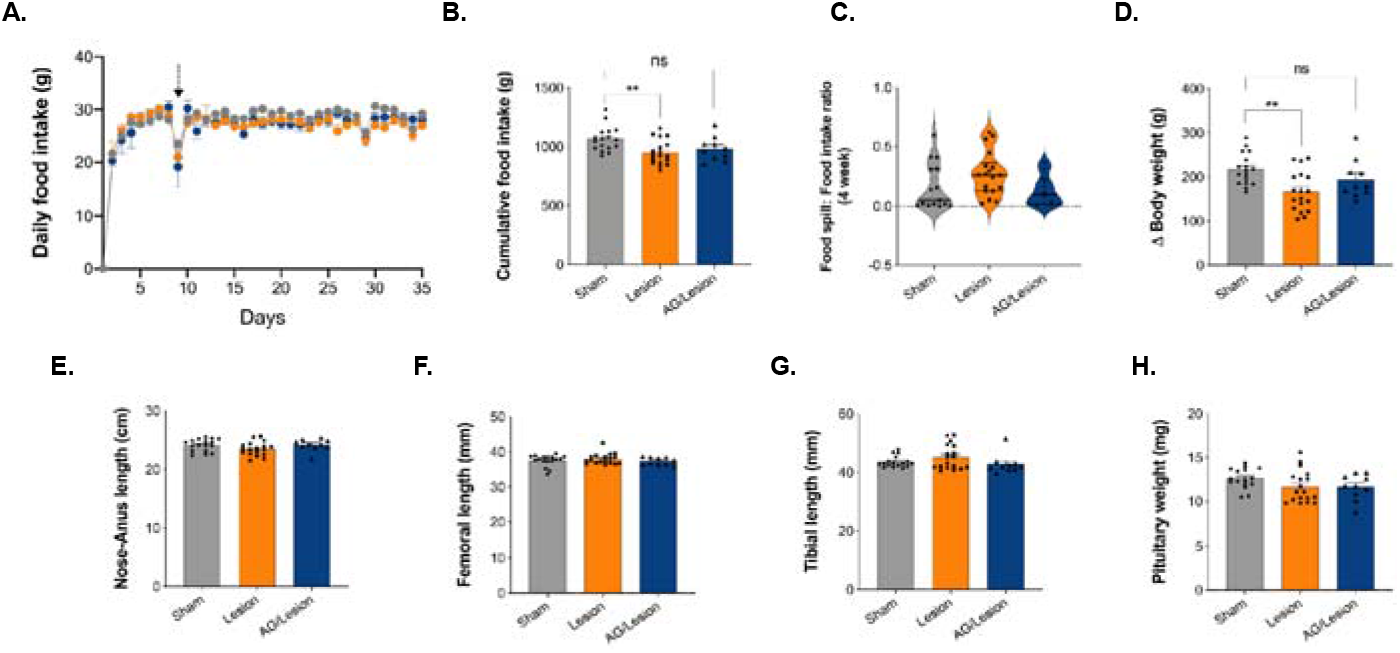
Acyl-ghrelin prevents weight loss induced by unilateral MFB 6-OHDA lesion. Daily food intake was unaffected (A), however, cumulative food intake over the 35 days of the study was significantly reduced in the Lesion (*P*<0.01) group (B). Acyl-ghrelin attenuated the 6-OHDA lesion-mediated increase in the Food spill:food intake ratio, although these changes were not statistically significant, as measured by Kruskal-Wallis test and Dunn’s post-hoc test as the data was not normally distributed (C). The 6-OHDA lesion-mediated reduction in body weight gain (*P*<0.01) was attenuated by acyl-ghrelin (D). There was no effect of treatment on any measure of skeletal growth, including nose-anus length (E), femoral length (F) and tibial length (G). Similarly, pituitary weight was not significantly affected by either treatment (H). Dotted line represents day of surgery. Statistical analysis by One-way ANOVA and Tukey post-hoc analysis. ** *P*<0.01 *vs* sham group. Data shown are mean ± SEM. (Sham, n = 17; Lesion, n = 19, Acyl-ghrelin/Lesion, n = 10 (AG = acyl-ghrelin)).

### Acyl-ghrelin treatment attenuated motor dysfunction induced by 6-OHDA lesion

Our use of a two (inner diet and outer spill) compartment food hopper for metabolic monitoring, enabled us to quantify the loss of the motor control of feeding. This revealed that by day 35, 4-weeks post-lesion, mean diet spillage in the unilateral 6-OHDA lesion group was 167% of that in the Sham group (Sham 0.1616 ± 0.04991 *vs* Lesion 0.2691 ± 0.04366, P=0.1124; Fig 1C). No loss of motor control was observed in the Acyl-ghrelin/Lesion group, where mean diet spillage was 80% of that in the Sham group (Sham 0.1616 ± 0.04991 *vs* Acyl-ghrelin/Lesion 0.1288 ± 0.04023; *P*>0.9999; Fig 1C). These data suggest that acyl-ghrelin pre-treatment may ameliorate broad motor impairments linked to feeding that were mediated by 6-OHDA lesion, possibly via preserved contralateral forelimb use (12). To determine whether acyl-ghrelin modulated motor function we utilised the commonly used amphetamine-induced rotation test to quantify the effect of unilateral lesion on motor asymmetry. The amphetamine-induced rotation reflects the imbalance in dopamine release between the denervated and intact striata and is widely used in the field to test neuroprotective interventions that preserve dopamine neurone function. On day 27, 3-weeks after 6-OHDA treatment, amphetamine triggered a significant increase in the number of complete ipsilateral rotations relative to sham-lesioned rats (Sham −1.065 ± 0.7906 *vs* Lesion 5.821 ± 1.132, *P*<0.01; Fig 2C). Notably, pre-treatment with acyl-ghrelin reduced the number of 6-OHDA mediated rotations by 74% to a level similar to the sham-lesioned rats (Sham −1.065 ± 0.7906 *vs* Acyl-Ghrelin/Lesion 1.519 ± 2.831, *P*>0.05; Fig 2C).

**Figure 2.**
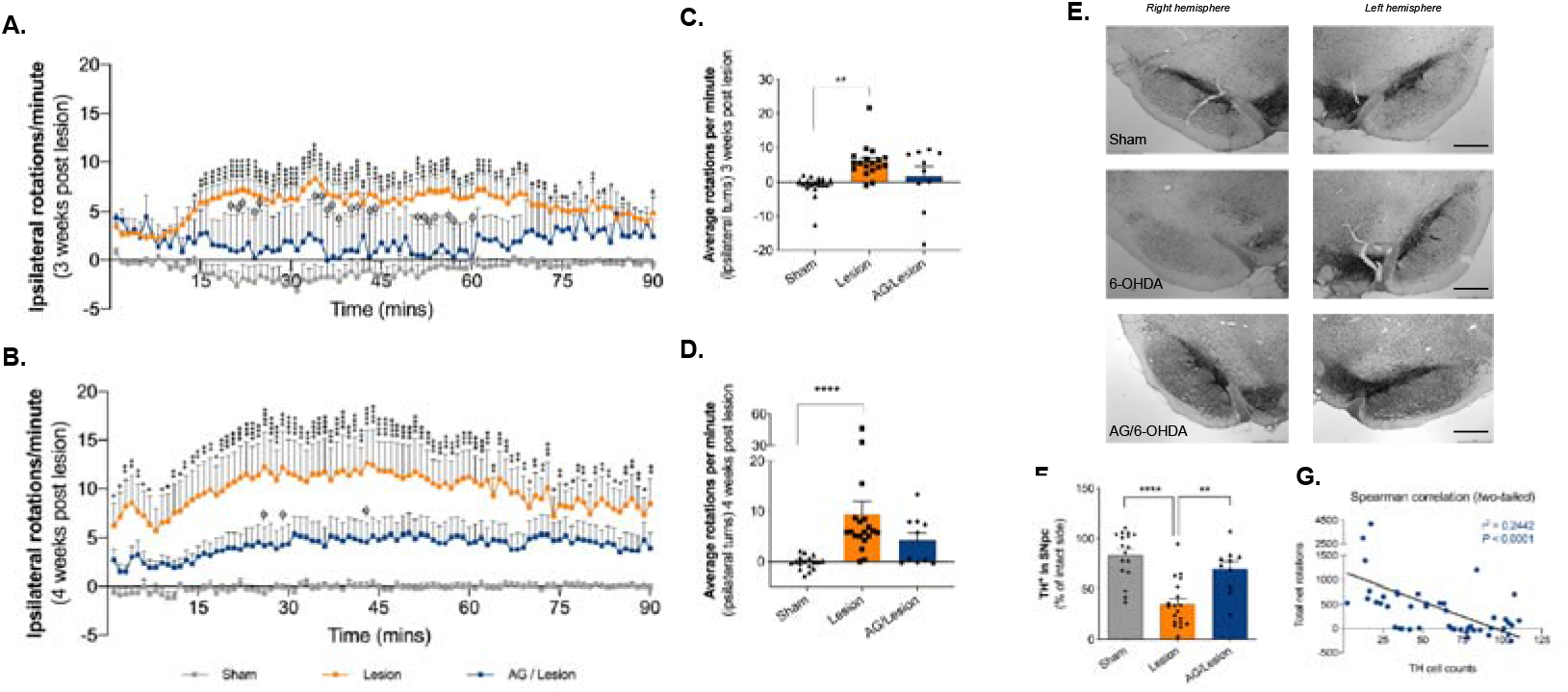
Acyl-ghrelin prevents motor and neurochemical deficits following unilateral MFB 6-OHDA lesion in adult rats. (A, B) The unilateral 6-OHDA lesion-mediated increase in amphetamine-induced rotations across the 90 minute testing period was attenuated by acyl-ghrelin pre-treatment 3- and 4-weeks post lesion (\**P*<0.05, \*\**P*<0.01, \*\*\**P*<0.001, \*\*\*\**P*<0.0001 *vs* sham group; □-*P*<0.05 *vs* lesion group). (C,D) Average rotations per minute were significantly increased by 6-OHDA at 3-weeks (*P*<0.01) and 4-weeks post lesion (*P*<0.0001), however, rotations were attenuated by acyl-ghrelin pre-treatment at both time points. (E) Photomicrographs of TH^+^ immunoreactive SNpc cells. (F) 6-OHDA decreased the number of TH^+^ SNpc cells by 58% (*P*<0.0001), whereas pre-treatment with acyl-ghrelin significantly reduced TH^+^ SNpc cell loss (*P*<0.01). (G) Spearman correlation (*two-tailed*) analysis of TH^+^ cell counts and total net rotations at 4 weeks. Statistical analysis was performed by Two-way (A,B) and One-way (C,D) ANOVA followed by Tukey’s post hoc test. (** *P*<0.01, **** *P*<0.001 *vs* lesion group). Data shown are mean ± SEM. Scale bar = 400um. (Sham, n = 17; Lesion, n = 19, AG/Lesion, n = 10 (AG = acyl-ghrelin)).

On day 35, rats that received the 6-OHDA toxin 4-weeks earlier had a more pronounced increase in the number of complete ipsilateral rotations relative to sham-lesioned rats (Sham −0.1670 ± 0.3211 *vs* Lesion 9.385 ± 2.615, *P*<0.0001; Fig 2D). As before, pre-treatment with acyl-ghrelin reduced the number of 6-OHDA mediated rotations to a level similar to the sham-lesioned rats (Sham −0.1670 ± 0.3211 *vs* Acyl-Ghrelin/Lesion 4.199 ± 1.54, *P*>0.05; Fig 2D). These data suggest that acyl-ghrelin pre-treatment prevented the motor deficits induced by 6-OHDA.

### Acyl-ghrelin treatment protected dopaminergic neurones from 6-OHDA-induced dysfunction

To determine whether the observed preservation of motor function corresponded with the number of dopaminergic cells within the midbrain, we quantified the number of TH ^+^ immunoreactive cells in the SNpc region of the midbrain in each experimental group. As predicted, the 6-OHDA lesion group had 58% less TH^+^ cells residing within the ipsilateral SNpc, relative to the sham-lesioned rats (Sham 83.64 ± 5.807 *vs* Lesion 34.96 ± 5.333, *P*<0.0001; Fig 2E,F). Notably, acyl-ghrelin pre-treatment inhibited the neurotoxic effect of lesion resulting in significantly more TH^+^ cells in the ipsilateral SNpc, relative to the 6-OHDA lesion group (Lesion 34.96 ± 5.333 *vs* Acyl-ghrelin/Lesion 70.15 ± 7.306, *P*<0.01; Fig 2E,F). The number of TH^+^ cells in the acyl-ghrelin pre-treatment group was not significantly different to the sham control group (Sham 83.64 ± 5.807 *vs* Acyl-ghrelin/Lesion 70.15 ± 7.306, *P*>0.05, Fig 2E,F). Consistent with this finding, increased TH^+^ SNpc cell number correlated with reduced amphetamine-induced ipsilateral rotations (Spearman *r*^2^ = 0.2442, *P*<0.0001; Fig 2G).

## Discussion

In this study, we investigated the effects of increasing circulating acyl-ghrelin for 7-days on long-term neurochemical and motor function in the unilateral MFB 6-OHDA lesion rat model of PD. Using the SNpc-dependent amphetamine-induced rotation task, 4-weeks after the toxin was administered, we evaluated motor performance. As expected, the 6-OHDA lesion group displayed significant rotational asymmetry compared to the sham-lesion group that is consistent with an imbalance in dopamine release between the denervated and intact striata (*P*<0.0001) (Fig 2A-D). However, elevating acyl-ghrelin levels for 7-days before introducing the toxin prevented a significant increase in ipsilateral rotations relative to the 6-OHDA lesion group (*P*<0.01) (Fig 2A-D). These data suggest that acyl-ghrelin treatment attenuated the loss of dopamine producing cells in the SNpc following 6-OHDA lesion and are in keeping with the finding that acyl-ghrelin administration ameliorates motor dysfunction in the mouse MPTP lesion model of PD (7) and the rat intra-striatal 6-OHDA model of PD (13). The major advantage of using this pre-clinical model is the quantifiable motor deficit induced by unilateral 6-OHDA lesion, and to the best of our knowledge, this is the first demonstration that acyl-ghrelin administration is efficacious in this rat model of PD.

Next, we performed histological analysis of dopamine (TH^+^) containing cells within the SNpc. The number of dopamine cells in the SNpc of the lesioned hemisphere was significantly reduced (58%) by 6-OHDA, relative to the non-lesioned hemisphere (*P*<0.0001) (Fig 2E,F). In contrast, acyl-ghrelin pre-treatment attenuated this loss, resulting in the preservation of significantly more dopamine cells in the lesioned SNpc, relative to the 6-OHDA lesion group (*P*<0.01) (Fig 2E,F). Indeed, there was no significant difference in the number of dopamine containing cells in the SNpc of the acyl-ghrelin treated and sham groups (*P*>0.05; Fig 2E,F). Consistent with the expectation of this pre-clinical model, the number of dopamine cells in the SNpc was inversely correlated with amphetamine-induced ipsilateral rotations (Spearman *r*^2^ = 0.2442, *P*<0.0001; Fig 2G). Our results are consistent with previously published findings that acyl-ghrelin prevents dopamine cell loss in pre-clinical toxin-based models of PD (7–10) and extend the repertoire of pre-clinical models used to validate the therapeutic potential of acyl-ghrelin signalling in PD.

In support of our hypothesis that acyl-ghrelin signalling may be a valid therapeutic target for neuroprotection in PD, evidence from humans PD patients suggests that post-prandial plasma total-ghrelin is reduced in PD individuals diagnosed with REM-sleep disorder, considered a prodrome of PD (14). Furthermore, fasting levels of total- and acyl-ghrelin were reduced in male and female subjects diagnosed with PD (15). Importantly, induced-pluripotent stem cell (iPSC)-derived dopaminergic neurones generated from patients carrying parkin gene (PARK2) mutations had significantly decreased GHS-R expression. Similarly, dopamine neurones of isogenic PARK2-iPSC lines mimicking PARK2 loss of function via CRISPR Cas9 technology also had reduced expression of GHS-R (16). More recently, we reported a reduction in the ratio of plasma acyl-ghrelin:unacyl-ghrelin specifically in individuals diagnosed with PD dementia when compared with cognitively intact PD and healthy control subjects (17). Given the limitations of administering putative therapeutic agents before the introduction of acute SNpc lesion, we suggest that studies using pre-clinical models linked with sporadic and familial PD are warranted to provide further insights into the role of ghrelin signalling in PD pathogenesis and progression. With growing interest in the role of systemic, particularly gastro-intestinal, factors in PD pathology (18), our findings provide further evidence that the stomach hormone, acyl-ghrelin, protects dopamine neurones in a pre-clinical model of PD. Indeed, given the efficacy of acyl-ghrelin in protecting dopaminergic neurone survival and function in the absence of the widely reported metabolic effects of hyperghrelinaemia, it is clear that human studies are warranted to test the efficacy of acyl-ghrelin peptide administration in slowing the progression of this debilitating disease.

